# High-Throughput Global Phosphoproteomic Profiling Using Phospho Heavy-Labeled-Spiketide FAIMS Stepped-CV DDA (pHASED)

**DOI:** 10.1101/2022.04.22.489124

**Authors:** Dilana E. Staudt, Heather C. Murray, David A. Skerrett-Byrne, Nathan D. Smith, Muhammad F. Jamaluddin, Richard G.S. Kahl, Ryan J. Duchatel, Zacary Germon, Tabitha McLachlan, Evangeline R. Jackson, Izac J. Findlay, Padraic S. Kearney, Abdul Mannan, Holly P. McEwen, Alicia M. Douglas, Brett Nixon, Nicole M. Verrills, Matthew D. Dun

**Author notes:** **CORRESPONDING AUTHOR INFORMATION:** Matthew Dun, School of Biomedical Sciences and Pharmacy, College of Health, Medicine and Wellbeing, University of Newcastle, NSW, 2308, Australia.

## Abstract

Global high-throughput profiling of oncogenic signaling pathways by phosphoproteomics is increasingly being applied to cancer specimens. Such quantitative unbiased phosphoproteomic profiling of cancer cells identifies oncogenic signaling cascades that drive disease initiation and progression; pathways that are often invisible to genomics sequencing strategies. Therefore, phosphoproteomic profiling has immense potential for informing individualized anti-cancer treatments. However, complicated and extensive sample preparation protocols, coupled with intricate chromatographic separation techniques that are necessary to achieve adequate phosphoproteomic depth, limits the clinical utility of these techniques. Traditionally, phosphoproteomics is performed using isobaric tagged based quantitation coupled with TiO_2_ enrichment and offline prefractionation prior to nLC-MS/MS. However, the use of isobaric tags and offline HPLC limits the applicability of phosphoproteomics for the analysis of individual patient samples in real-time. To address these limitations, here we have optimized a new protocol, <u>p</u>hospho-<u>H</u>eavy-l<u>a</u>beled-spiketide FAIM<u>S</u> St<u>e</u>pped-CV <u>D</u>DA (pHASED). pHASED maintained phosphoproteomic coverage yet decreased sample preparation time and complexity by eliminating the variability associated with offline prefractionation. pHASED employed online phosphoproteome deconvolution using high-field asymmetric waveform ion mobility spectrometry (FAIMS) and internal phosphopeptide standards to provide accurate label-free quantitation data. Compared with our traditional tandem mass tag (TMT) phosphoproteomics workflow and optimized using isogenic FLT3-mutant acute myeloid leukemia (AML) cell line models (n=18/workflow), pHASED halved total sample preparation, and running time (TMT=10 days, pHASED=5 days) and doubled the depth of phosphoproteomic coverage in real-time (phosphopeptides = 7,694 pHASED, 3,861 TMT). pHASED coupled with bioinformatic analysis predicted differential activation of the DNA damage and repair ATM signaling pathway in sorafenib-resistant AML cell line models, uncovering a potential therapeutic opportunity that was validated using cytotoxicity assays. Herein, we optimized a rapid, reproducible, and flexible protocol for the characterization of complex cancer phosphoproteomes in real-time, highlighting the potential for phosphoproteomics to aid in the improvement of clinical treatment strategies.

## INTRODUCTION

Mass spectrometry approaches for global high-throughput quantitation of cellular phosphoproteomes have been increasingly applied to cancer specimens as they provide powerful tools for the identification of signaling pathways including kinases, phosphatases and cell cycle regulators that drive disease initiation and progression (1–3). Deregulation of kinase and phosphatase activity plays a critical role in cancer development and relapse (4–7), highlighting kinases as important therapeutic targets in the clinic (8, 9). This is particularly the case for FLT3 kinase-driven acute myeloid leukemia (AML) patients. The FLT3 receptor tyrosine kinase is recurrently mutated in AML patients and is a target for FLT3 inhibitor therapy. The most common mutations are internal tandem duplications (ITD) and kinase domain mutations (e.g. D835). Resistance to FLT3 inhibitor therapy is often associated with the emergence of dual FLT3-ITD/D835 mutations, however the pathways mediating drug resistance are yet to be fully characterized (3, 5, 6). Therefore, phosphoproteomic profiling of the activated kinases responsible for driving downstream oncogenic signaling cascades using cancer patients’ specimens in real-time, provides an opportunity to repurpose clinically relevant therapeutic interventions (10–14), and thus aid in the development of individualized treatment strategies that may improve overall survival.

Several methods have been developed for the quantitative characterization of phosphoproteins in complex biological samples using shotgun proteomics (15–18). Stable isotope-labeling strategies such as tandem mass tag (TMT) approaches have become increasingly popular due to the capability to multiplex analysis of up to 18 complex matrices simultaneously. TMT protocols enable samples to be pooled prior to nano liquid chromatography–tandem mass spectrometry (nLC-MS/MS) therefore saving instrument time and reducing technical variations in the workflow. However, the highly complex nature of cancer phosphoproteomes necessitates that TMT protocols are coupled with phosphopeptide enrichment and sample pre-fractionation prior to nLC-MS/MS analysis in order to achieve adequate phosphoproteome resolution. Additionally, the high cost of reagents, fixed number of samples, and sample preparation time and complexity combine to limit the utility of TMT protocols for the ad hoc assessment of patient specimens in real-time.

Label-free quantitation (LFQ) strategies provide quantitative phosphoproteomic data without the use of isotopic-tags, mainly through the direct inference of protein abundance using the measured intensity of detected peptides, or indirect inference based on the number of phosphopeptide-spectrum matches (PSMs) obtained for each protein (19). LFQ protocols have the capacity to overcome some of the TMT-workflow limitations by reducing the complexity of sample preparation, saving both time and on costly reagents. Additionally, there is no limit to the number of matrices to be analyzed, thus enabling the comparison of larger sets of samples than when using label-based approaches. Such strategies therefore hold obvious appeal in the context of highly aggressive forms of cancer in which the design of appropriate treatment strategies is time-sensitive, and hence the ability to rapidly perform phosphoproteomic profiling on a high number of samples is of critical importance. However, label-free strategies have their own limitations, which include the inherent variability of individual sample preparation and loading, and the requisite number of replicates. Additionally, chromatographic conditions and the semirandom nature of data acquisition also have an impact on sample reproducibility (20). The addition (spike-in) of known concentrations of standard heavy-labeled exogenous phosphopeptides for sample normalization however, can help to overcome some of these limitations (16, 21), and therefore provides a strategy to normalize protein expression and phosphorylation abundance from different cancer specimens analyzed at any time. Additionally, by interfacing phosphopeptide enrichment with separation via FAIMS prior to high-resolution mass spectrometry (MS) the collection of single-shot proteomic data is possible without the need to perform conventional two-dimensional liquid chromatography (2D-LC) approaches (22), and provides deep phosphoproteomic coverage to identify cancer-associated drug targets, in real-time.

In seeking to combine the salient features of these analytical modalities, here we report the optimization of a new protocol that employs online phosphoproteome deconvolution in tandem with LFQ in the presence of internal control heavy-labeled standards. This protocol was developed to identify kinases driving disease progression and therapy resistance in real-time. To determine the pre-clinical utility of this approach, pHASED was applied to isogenic FLT3-mutant AML cell lines resistant to the tyrosine kinase inhibitor sorafenib.

## EXPERIMENTAL PROCEDURES

### Cell Culture

Murine hematopoietic progenitor FDC-P1 cells were stably transduced with either human wildtype (WT) *FLT3*, *FLT3*-ITD, *FLT3*-D835Y, *FLT3*-D835V, *FLT3*-ITD/D835V, or *FLT3*-ITD/D835Y by retroviral transduction (6), confirmed by standard Sanger sequencing (Suppl Materials and Methods). FDC-P1 FLT3-transduced lines were maintained in standard culture conditions (5% CO_2_, 37°C) in DMEM medium (Thermo Fisher Scientific) with the addition of 10% FBS, and 20mM HEPES (N-2-hydroxyethylpiperazine-N’-2-ethanesulfonic acid). A total of 50 ng/mL human FLT3-ligand (Biolegend) was added to FLT3-WT cells, whereas FLT3-mutant lines are factor-independent and were therefore maintained in growth factor free media. All cell lines were routinely confirmed to be free of mycoplasma contamination using a MycoAlert mycoplasma detection kit (Lonza; Basel, Switzerland).

### Sample Preparation and Protein Extraction

#### TMT-based phosphopeptide quantification

Snap frozen transduced FDC-P1 cells expressing human wildtype-*FLT3* and AML associated FLT3-mutations were lysed in 100 μL of ice-cold 0.1 M Na_2_CO_3_, pH 11.3 containing protease and phosphatase inhibitors (Sigma, cat. #P8340-5ML, and #4906837001 respectively), by sonication (2 × 20 s cycles, 100% output power) (as described (23–25)). Protein concentration was determined using a Bicinchoninic acid (BCA) protein estimation assay, as per manufacturer’s instructions (Thermo Fisher Scientific). Protein samples were then diluted in 6 M Urea/2 M Thiourea and reduced using 10 mM dithiothreitol (DTT) by incubation for 30 min at room temperature (RT). Reduced cysteine residues were then alkylated using 20 mM iodoacetamide by incubation for 30 min at RT in the dark. Enzymatic digestion was achieved using Trypsin/Lys-C mixture (Promega) at an enzyme-to-substrate ratio of 1:50 (w/w) and incubated for 3 h at RT. Triethylammonium bicarbonate (TEAB, 50 mM, pH 7.8) was then added to dilute urea concentration below 1 M, and samples were incubated overnight at RT. Lipid precipitation was performed using formic acid and trichloroacetic acid (TCA). Briefly, a final concentration of 2% formic acid was added to each sample, prior to centrifugation at 14,000 g for 10 min. Remaining lipopeptides were then precipitated with 20% (w/w) TCA and incubated on ice for at least 1 h prior to centrifugation. Pellets were washed with ice cold 0.01 M hydrochloric acid (HCl)/90% acetone and supernatants containing peptides were combined. Peptides were desalted using Oasis HLB solid phase extraction (SPE) cartridges and a VisiprepTM SPE Vacuum Manifold (12-port model; Sigma). The SPE cartridges were activated using 100% acetonitrile (ACN) and equilibrated using 0.1% trifluoroacetic acid (TFA). Acidified samples (pH < 3) were loaded onto SPE cartridges with liquid passed through the solid phase dropwise using vacuum pressure. The cartridges were washed with 0.1% TFA followed by sequential elution of peptides using 60% ACN/0.1% TFA and 80% ACN/0.1% TFA. Eluted peptides were then resuspended in TEAB (50 mM, pH 8) and quantitated using a peptide fluorescence assay kit (Thermo Fisher Scientific). 100 µg from each of the samples were individually labeled using tandem mass tags (Supplemental Table S1; TMT-10plex 3 × kits, Thermo Fisher Scientific, Bremen DE, Germany) and mixed at a 1:1 ratio. Phosphopeptides were isolated from the proteome using titanium dioxide (TiO_2_) as previously described (11) before offline hydrophilic interaction liquid chromatography (HILIC) using a Dionex Ultimate 3000RSLC nanoflow HPLC System (Thermo Fisher Scientific).

#### pHASED

peptide preparation was performed the same as for TMT and peptides desalted as above. Following activation and equilibration, SPE cartridges were blocked with 33 µg of trypsin-digested bovine serum albumin (BSA) peptides prior to sample clean-up. Peptides were sequentially eluted using 60% ACN/0.1% TFA, and 80% ACN/0.1% TFA, and the eluates were quantified using a Qubit 2.0 Fluorometer, as per manufacturer’s instructions (Thermo Fisher Scientific). A total of 200 µg of peptide per sample was utilized for TiO_2_ enrichment. Spike-in heavy-labeled phosphorylated peptides (Supplemental Table S2; including individually tyrosine, threonine or serine phosphorylated heavy-labeled spiketides, 8 fmol/200 μg of sample) were added as internal controls. Phosphopeptide enrichment was modified based on previous protocols (11, 13, 18). In brief, each peptide sample was suspended in 80% ACN, 5% TFA, and 1 M glycolic acid (loading buffer). TiO_2_ beads were added at 0.6 mg per 100 µg peptide (w/w), and samples were mixed at RT for 15 min. The supernatant was incubated with half the amount of fresh TiO_2_ beads, and resultant supernatants containing non-phosphorylated peptides (non-modified = NM fraction) were removed and stored. The two sets of beads with bound phosphopeptides were pooled using 100 μL of loading buffer, followed by sequential washing with 80% ACN/1% TFA, and 10% ACN/0.1% TFA. Phosphopeptides were eluted with 28% ammonia hydroxide solution (1% v/v, pH 11.3) then passed through a C8 stage tip to remove residual beads (18). Phosphopeptides were lyophilized completely prior to resuspension in 2% ACN/0.1% TFA for nLC-MS/MS analysis.

### Nanoflow Liquid Chromatography Tandem Mass Spectrometry Mass Spectrometry (nLC-MS/MS)

#### TMT-based phosphopeptide quantification

LC tandem mass spectrometry (MS/MS) was performed on 9 phosphopeptide enriched HILIC fractions using a Q-Exactive Plus hybrid quadrupole-Orbitrap MS system (Thermo Fisher Scientific) coupled to a Dionex Ultimate 3000RSLC nanoflow HPLC system (Thermo Fisher Scientific). Approximately 700 ng of phosphopeptide per HILIC fraction were loaded onto an Acclaim PepMap100 C18 75 μm × 20 mm trap column (Thermo Fisher Scientific) for pre-concentration and online desalting. Separation was then achieved using an EASY-Spray PepMap C18 75 µm × 25 cm column (Thermo Fisher Scientific) employing a linear gradient from 5 to 35% acetonitrile at 300 nL/min over 127 min. The Q-Exactive Plus MS System (Thermo Fisher Scientific) was operated in full MS/data-dependent acquisition MS/MS mode (DDA). The Orbitrap mass analyzer was used at a resolution of 70,000, to acquire full MS with an m/z range of 380–2000, incorporating a target automatic gain control value of 1e^6^ and maximum fill times of 50 ms. The 20 most intense multiply charged precursors were selected for higher-energy collision dissociation (HCD) fragmentation with a normalized collisional energy of 32. MS/MS fragments were measured at an Orbitrap resolution of 35,000 using an automatic gain control target of 5e^5^ and maximum fill times of 120 ms.

#### pHASED

reverse phase nanoflow LC-MS/MS was performed using a Dionex Ultimate 3000RSLC nanoflow high-performance liquid chromatography system coupled with an Orbitrap Exploris 480 MS equipped with a front-end FAIMS Interface (Thermo Fisher Scientific). Approximately 700 ng of phosphopeptide per CV were loaded onto an Acclaim PepMap 100 C18 75 μm × 20 mm trap column for pre-concentration and online de-salting. Separation was then achieved using an EASY-Spray PepMap C18 75 μm × 25 cm, employing a gradient of 0-35% solvent B (solvent A = 0.1% formic acid, solvent B = 90% ACN, 0.1% formic acid) at a flow rate of 250 nL/min over 75 min. The mass spectrometer was operated in positive mode with the FAIMS Pro interface. Four compensation voltages (CV; −70, −60, −50, −40) were individually run for each biological triplicate. Full MS/data dependent acquisition (DDA) was performed using the following parameters: Orbitrap mass analyzer set at a resolution of 60,000, to acquire full MS with an m/z range of 350-1200, incorporating a standard automatic gain control target of 1e^6^ and maximum injection time of 50 ms. The 20 most intense multiply charged precursors were selected for higher-energy HCD with a collisional energy of 30. MS/MS fragments were measured at an Orbitrap resolution of 15,000 incorporating a normalized automatic gain control target of 250% and a maximum injection time of 120 ms.

### Data Processing and Bioinformatic Analysis

Data analysis was performed using Proteome Discoverer (Thermo Fisher Scientific). Sequest HT was used to search against UniProt *Mus musculus* database (25,280 sequences, downloaded 30/05/20 for TMT; 17,462 sequences, downloaded 23/03/21 for pHASED) and *Homo sapiens* FLT3 FASTA file containing WT and mutant FLT3 sequences (3 sequences, downloaded 21/02/20 in both experiments). Database searching parameters included up to two missed cleavages, precursor mass tolerance set to 10 ppm and fragment mass tolerance of 0.02 Da. Cysteine carbamidomethylation was set as a fixed modification while dynamic modifications included oxidation (M), phosphorylation (S, T, Y), acetylation (K), methylation (K) and deamidation (N, Q). In addition, N-terminus TMT6plex was set as fixed modifications for TMT-labeled samples. Interrogation of the database was performed to evaluate the false discovery rate (FDR) of peptide identification based on q-values estimated from the target-decoy search approach using Percolator. An FDR rate of 1% was set at the peptide level to filter out target peptide spectrum matches over the decoy-peptide spectrum matches. Additionally for pHASED samples, heavy-labeled 13C(6)15N(2) (K), and 13C(6)15N(4) (R) modifications were included as dynamic modifications to identify spiked-in heavy-labeled phospho-spiketides. To account for variations in sample injection, reporter ion abundances were normalized to total peptide amount for the TMT-labeled protocol, and the spiked-in heavy-labeled phosphopeptides included as FASTA file for pHASED (Suppl Fig. S1). For quantification and comparison, each ratio was transformed to log2 scale (log2 ratio).

### Experimental Design and Statistical Rationale

Phosphoproteomic data analysis was performed using six FDC-P1 isogenic cell lines (n=3 biological replicates). Four compensation voltages (CV; −70, −60, −50, −40) were individually analyzed for each biological replicate. Differentially expressed phosphopeptides and phosphorylation sites were defined as those with a significant (*p*≤0.05) log2 fold change ≥0.25 or ≤−0.25. Differences between sample groups were analyzed by unpaired Student’s t-tests or one-way ANOVA and considered significant when *p*≤0.05. Graphical data was analyzed and prepared using Perseus (1.6.2.2), String (11.5), CytoScape (3.9.1), and GraphPad Prism (9.0.1). Results are presented as mean values ± SEM.

### Ingenuity Pathway Analysis

Ingenuity Pathway Analysis software (IPA; Qiagen) was used to analyze each phosphoproteomic dataset (as previously described (11, 13)). Canonical pathways, upstream regulators, and disease and function analyses were generated and assessed based on *p*-value.

### Kinase-Substrate Enrichment Analysis

Kinase-Substrate Enrichment Analysis (KSEA App, version 1.0) (26) was used to analyze phosphorylated sites based on PhosphoSitePlus (27) kinase-substrate dataset, and a *p* ≤0.05 cut-off.

### Cytotoxicity Assays

Cell lines were treated with the FLT3 inhibitor sorafenib (Selleckchem) (28), and ATM inhibitor KU-60019 (Selleckchem) (29) either alone or in combination. Cells were seeded into 96 well plates at 2e^4^ cells per well, and viability following treatments was measured using Resazurin (excitation 544 nm, emission 590 nm; 0.6 mM Resazurin, 78 μM Methylene Blue, 1 mM potassium hexacyanoferrate (III), 1 mM potassium hexacyanoferrate (II) trihydrate (Sigma), dissolved in sterile phosphate buffered saline). Synergy of dose-response and combined effect of the two drugs were assessed using the method of Bliss independence model (30).

## RESULTS

### pHASED reduced sample preparation time by half whilst providing improved phosphopeptide quantification compared to the TMT workflow

The new label-free phosphoproteomic enrichment and MS protocol ‘pHASED’ described herein, couples phosphopeptide enrichment strategies optimized by Engholm-Keller et.al., (2012) (18) and LFQ using heavy-labeled internal phospho-spiketides and FAIMS interface optimized by Alexander et.al., (2018) (22), to decrease sample preparation time and increase phosphoproteome deconvolution and coverage for the analysis of samples in real-time (Fig. 1). We performed initial comparison of our optimized pHASED with traditional TMT phosphoproteomic protocols using six isogenic cell line models of FLT3-mutant AML in biological triplicate (Table 1; n = 36 samples). The sample preparation in pHASED saves time due to the substitution of TMT-labeling with the spike-in of heavy-labeled phospho-spiketides of known concentration in order to normalize sample injection and phosphopeptide quantitation. We replaced offline HILIC for online deconvolution using FAIMS interface employing external stepping of four different compensation voltages (CV; −70, −60, −50, −40) over a 75 min gradient. Individual sample injection per CV provided more flexibility to the experiment, however, increased LC-MS/MS time by 1.4 days (2 days TMT; 4 days pHASED). This longer instrument time however, is compensated by the reduction of sample preparation time by half, requiring an overall ∼5 days for completion of pHASED experiment, whereas ∼10 days are required for TMT (Fig. 1*A*, 1*B*).

**Figure 1:**
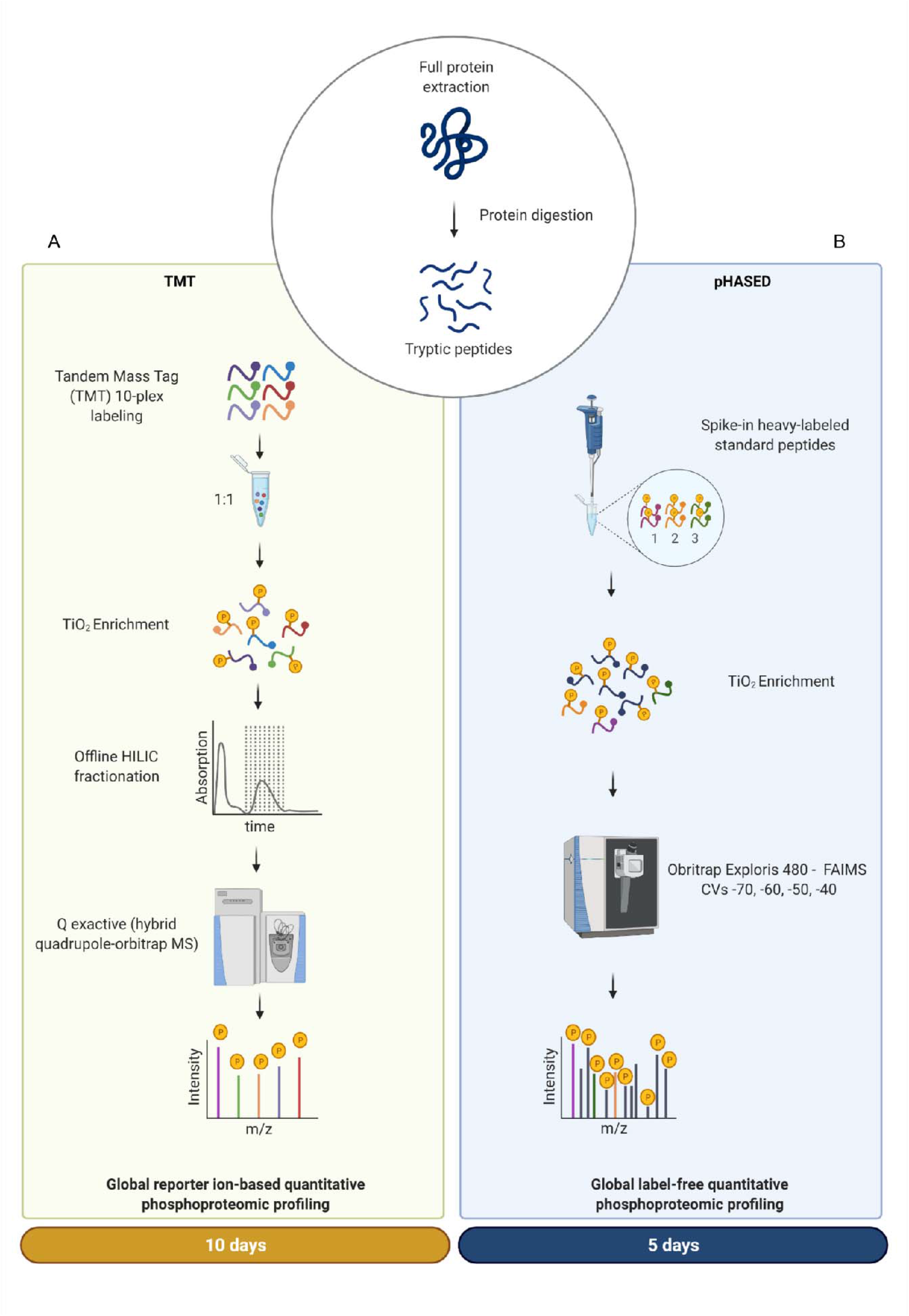
Overview of sample preparation and instrument workflow to compare quantitative phosphoproteomics using TMT and pHASED. Proteins were extracted from target cell lines and digested into peptides. *A)* One hundred micrograms of peptide/sample was labeled with TMT 10-plex isobaric tags, mixed 1:1, and enriched for phosphopeptides prior to offline HILIC fractionation and analysis on a Q Exactive Orbitrap MS. *B)* In the optimized pHASED workflow, two hundred micrograms of digested peptides were separated for enrichment. Known concentrations of spike-in heavy labeled phosphopeptides were added to each sample prior to phosphopeptide enrichment, and enriched phosphopeptides were then injected into an Orbitrap Exploris 480 coupled with a FAIMS interface using four different compensation voltages (CVs; −70V, −60V, −50V and −40V). Figure created with BioRender.com.

**Table 1:** Isogenic cellular models of FLT3 mutant AML analyzed by TMT and pHASED protocols in biological triplicate ((n=36/technique)

| Cell line | Species | Biological replicates | Samples | Model of AML stage | Sorafenib sensitivity |
| --- | --- | --- | --- | --- | --- |
| FDC-P1 | Mouse | 3 | FLT3wt | Normal | Sensitive |
| FDC-P1 | Mouse | 3 | FLT3-ITD | Diagnosis | Sensitive |
| FDC-P1 | Mouse | 3 | FLT3-D835V | Diagnosis | Sensitive |
| FDC-P1 | Mouse | 3 | FLT3-D835Y | Diagnosis | Sensitive |
| FDC-P1 | Mouse | 3 | FLT3-ITD/D835V | Relapse | Resistant |
| FDC-P1 | Mouse | 3 | FLT3-ITD/D835Y | Relapse | Resistant |

To determine the utility of each protocol, we examined the PSMs of each experiment (Fig. 2). Analysis of the charge states (Fig. 2*A*) and precursor ion mass-to-charge ratios (m/z) (Fig. 2*B*) for the two protocols demonstrated that our traditional TMT approach identified a higher percentage of +2 and +3 charged precursor ions, ranging between 400-700 m/z, whereas pHASED identified a greater number of +4, +5, and +6 precursors and higher m/z ratios (700-1200). The fractionation profile of the four CVs applied in pHASED were analyzed by comparing the PSMs acquired in each CV (Fig. 2*C*-*F*). More unique PSMs were identified in lower CVs, such as −70V and −60V compared to −50V and −40V (Fig. 2*C*). Interestingly, similar FAIMS distributions were seen for phosphopeptides as previously reported for studies employing FAIMS to characterize the non-modified proteome (22). For most charge states, the lower the CV the lower the average m/z (Fig. 2*D*, 2*E*), except for +6 charged precursors, which showed similar average m/z across CVs (Fig. 2*E*). Analysis of the overlapping distributions of common and unique phosphoproteins identified showed deep phosphoproteome coverage across all four CVs (Fig. 2*F*) (31).

**Figure 2.**
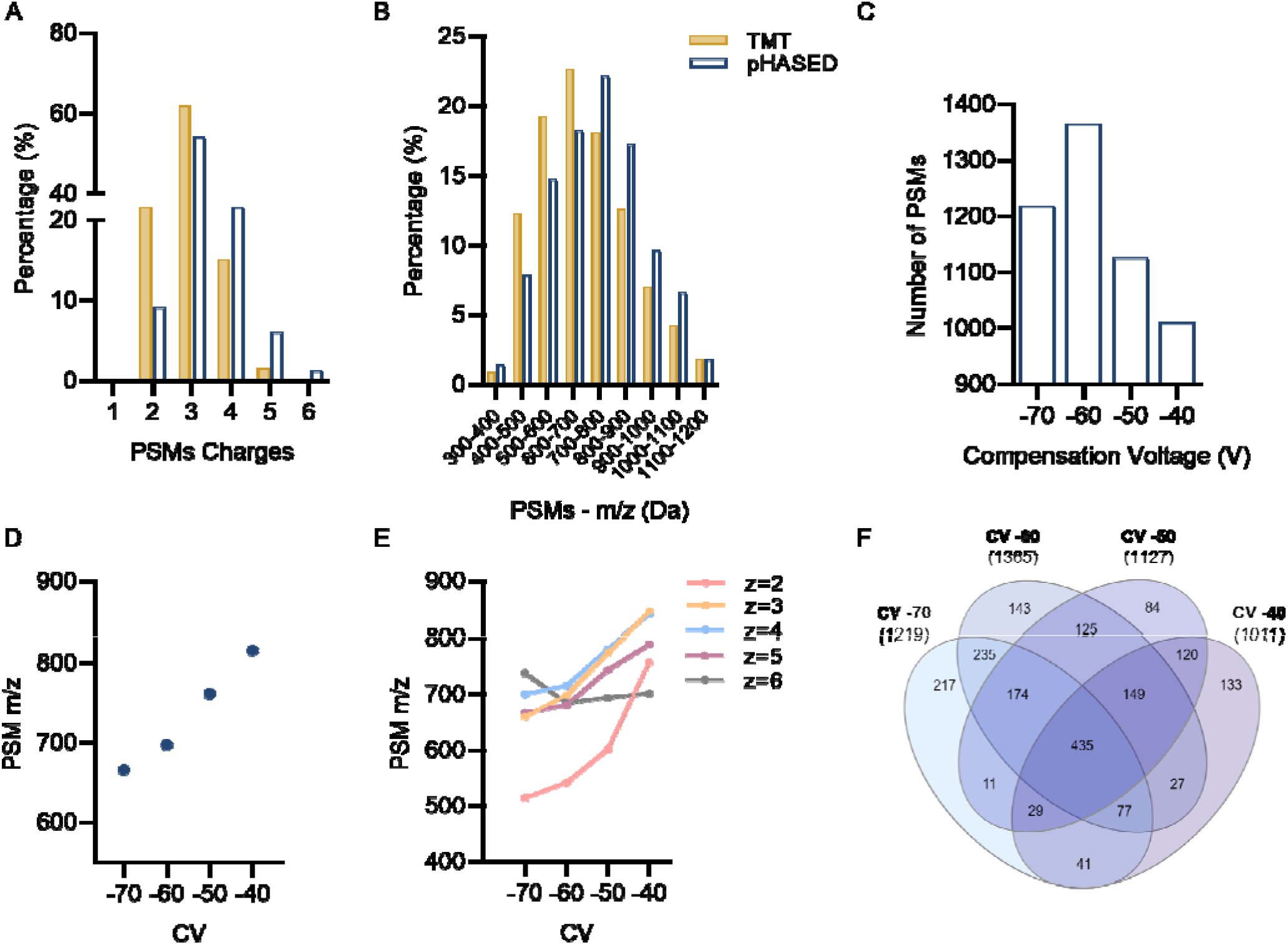
Acquisition profile of phosphopeptide-spectrum matches (PSMs) resulting from TMT compared to pHASED. *A)* Percentage of charge states of all peptide ions selected for MS/MS in TMT and pHASED experiments*. B)* Distribution of precursor ions identified in each experiment, stratified according to m/z. *C*) Percentage of PSMs identified in each CV. *D*) Average m/z of all PSM features acquired in each CV. *E*) Average m/z of PSM features for charge states acquired in each CV. *F*) Venn distribution of unique phosphoprotein accessions identified in CVs −70V, −60V, −50V and −40V shows overall coverage of common and unique acquisitions detected in each CV. Venn diagram created with InteractiVenn.

To examine the reproducibility of the quantification achieved using TMT and pHASED we performed Pearson Correlation analysis of the biological replicates (n=3) across each cell line (n=6) (Fig. 3). Correlation was performed by plotting normalized phosphopeptide abundances from each biological replicate per sample in a correlation matrix. This analysis revealed increased quantification reproducibility in all biological replicates of samples analyzed with pHASED (Fig. 3*A,* 3*C*) in comparison to the TMT protocol (Fig. 3*B,* 3*C*), which presented a moderate correlation between replicates and samples. These results indicate that, in our hands, pHASED performed as a more consistent phosphopeptide quantification tool, and therefore may yield more biologically relevant data.

**Figure 3.**
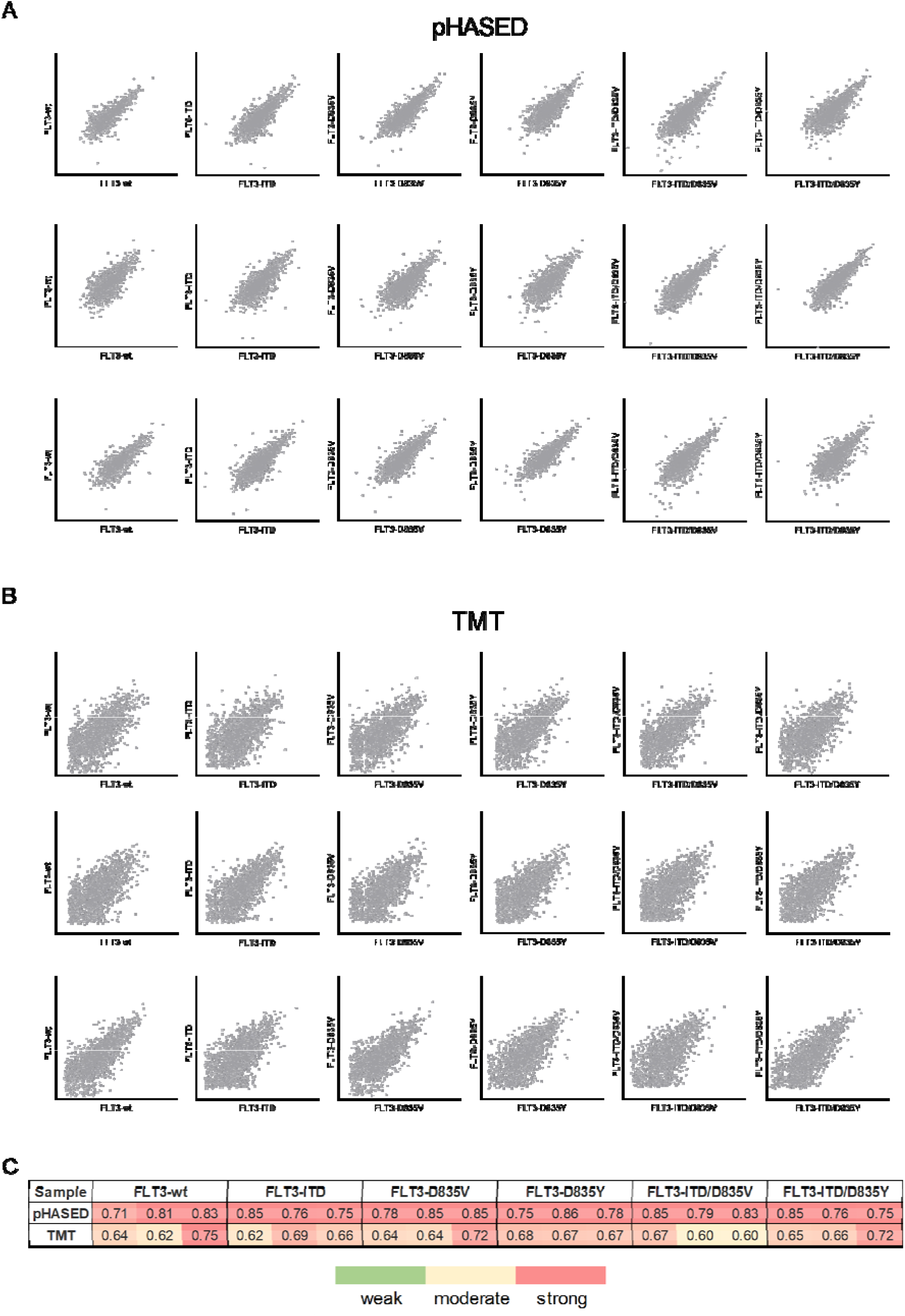
Quantification reproducibility between biological replicates TMT and pHASED experiments. Pearson correlation profiles for biological replicates (n=3) of six isogenic models of acute myeloid leukemia analyzed by *A*) TMT, and *B*) pHASED label-free experiments. *C*) Heatmap comparison of all three correlation scores achieved by from isogenic cell line for TMT and pHASED. Correlation was performed using normalized abundances in Perseus, and graphs were plotted using GraphPad Prism 8.4.3.

### pHASED provided in-depth phosphoproteome coverage

Our TMT approach identified 1,958 phosphoproteins and 3,861 phosphorylated peptides (FDR 1%), whereas a total of 1,587 phosphoproteins and 7,694 phosphorylated peptides were identified using pHASED (FDR 1%) (Fig. 4*A*, Supplemental Tables S*3*, S*4*). Both protocols identified similar S:T:Y ratios, in accordance with previous findings reporting a phosphorylation ratio of 86:12:2 (%) (32). The overall success of phosphopeptide enrichment for each experiment was 72% for TMT, and 93% for pHASED (Fig. 4*A*), indicating good enrichment efficiency in both experiments. Furthermore, pHASED identified an overall higher number of multi-phosphorylated peptides (2,800 singly-, 1,348 doubly-, and 323 triply-phosphorylated peptides, FDR 1%) compared to our traditional TMT approach (2,069 singly-, 334 doubly-, and 11 triply-phosphorylated peptides, FDR 1%) (Fig. 4*B*). Notably, overall increased phosphoprotein coverage and identification of more peptides per protein was achieved by pHASED (Fig. 4*C*, 4*D*). Comparison between identified phosphoprotein accessions across all datasets showed a 47% overlap between pHASED and TMT (Fig. 4*E*, and Supplemental Table S*5)*.

**Figure 4.**
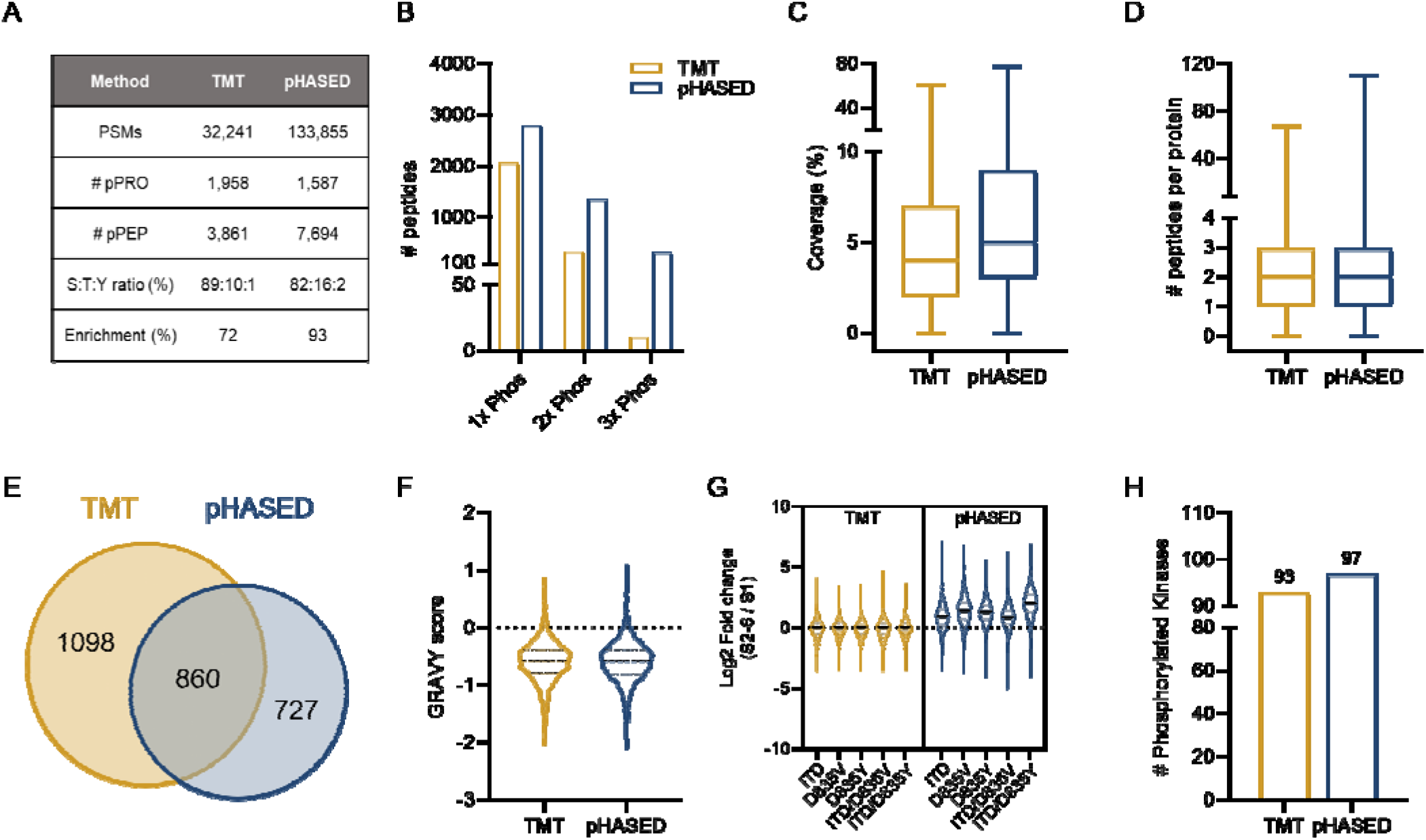
Analysis of phosphoproteome coverage and phosphopeptide characteristics identified using TMT compared with pHASED. *A)* Summary of MS acquisition comparing TMT and pHASED experiments. *B)* Number of single and multi-phosphorylated peptides identified in each experiment comparing TMT and pHASED. *C*) Phosphoprotein coverage comparing TMT and pHASED experiments. *D*) Number of phosphopeptides per protein identified in TMT and pHASED experiments. *E*) Overlap of phosphoprotein accessions comparing TMT and pHASED experiments. *F)* GRAVY score of peptides identified in each TMT and pHASED experiment irrespective of PTMs. *G)* Median distribution of log2 ratios for phosphopeptide changes in FLT3 mutants (S2-S6) compared to FLT3-wt (S1) in TMT and pHASED experiments. *H*) Number of phosphorylated master protein kinases identified in each TMT and pHASED experiment (*p*<0.01).

Both pHASED and TMT identified similar numbers of hydrophobic peptides, with both protocols preferencing the identification of hydrophilic phosphopeptides (Fig. 4*F*). For quantification, normalization of TMT samples was achieved based on total peptide amount per TMT channel, whereas each sample analyzed by pHASED was normalized with spike-in phospho-spiketides of known concentration. Measured abundance ratios were then transformed to log2 scale (log2 ratio). The distribution of phosphopeptide log2 fold-changes comparing FLT3 mutant sample to FLT3-wt cell lines measured by pHASED showed a greater dynamic range than those measured by TMT-based quantification (Fig. 4*G*), with a mean of log2 fold-change closer to 0 for all cell lines in the TMT approach compared to pHASED (mean log2 1.29; *p*=0.0002).

### pHASED identified relevant therapeutic drug targets in drug resistant AML

pHASED and TMT identified similar numbers of phosphoproteins with kinase activity (Fig. 4*H; 1% FDR*). In accordance with our previous findings of divergencies between the number of identified phosphoproteins in TMT compared to pHASED (Fig. 4*E*), 39% of kinases were identified to be common to both analyses (Supplemental Tables S*6-8*). Despite differences, the numbers of kinases identified showed that both protocols were effective for the identification of clinically relevant drug targets that could potentially aid in the design of treatment strategies for cancer patients. Analysis of the phosphorylation profile of kinases identified in FLT3-ITD, resistant FLT3-ITD/D835V, and resistant FLT3-ITD/D835Y cell lines was performed to investigate the clinical utility of pHASED, and results were compared to our TMT approach (Fig. 5). FLT3-ITD mutations are seen in approximately 27% of AML patients at diagnosis and are associated with a high risk of relapse (3). Resistance to commonly used FLT3 inhibitors including sorafenib (33–39) (Supplemental Table S*9*) occurs following FLT3-ITD+ AML cells acquiring a secondary point mutation in the kinase domain of FLT3 (FLT3-ITD/D835V and FLT3-ITD/D835Y, henceforth referred to as “double mutant”). Cytotoxicity assays using the FLT3 inhibitor sorafenib confirmed the resistant phenotype of FLT3-ITD/D835V and FLT3-ITD/D835Y mutants (Fig. 5*A*), which presented an average 47-fold increase in sorafenib IC_50_ in comparison to FLT3-ITD cell lines (IC_50_ 4.2 µM, 2.7 µM, 0.073 µM, FLT3-ITD/D835V, FLT3-ITD/D835Y and FLT3-ITD respectively). pHASED identified a significantly increased number of kinases that showed differential phosphorylation in both resistant cell lines in comparison to the TMT approach, particularly in FLT3-ITD/D835Y mutants (Fig 5*B; p*=0.0009). In addition, analysis of kinases showing significantly altered phosphorylation in double mutants compared with FLT3-ITD cells (log2 +/- 0.25; *p*≤0.05) showed greater dynamic range via pHASED compared to TMT (Fig. 5*C*).

**Figure 5.**
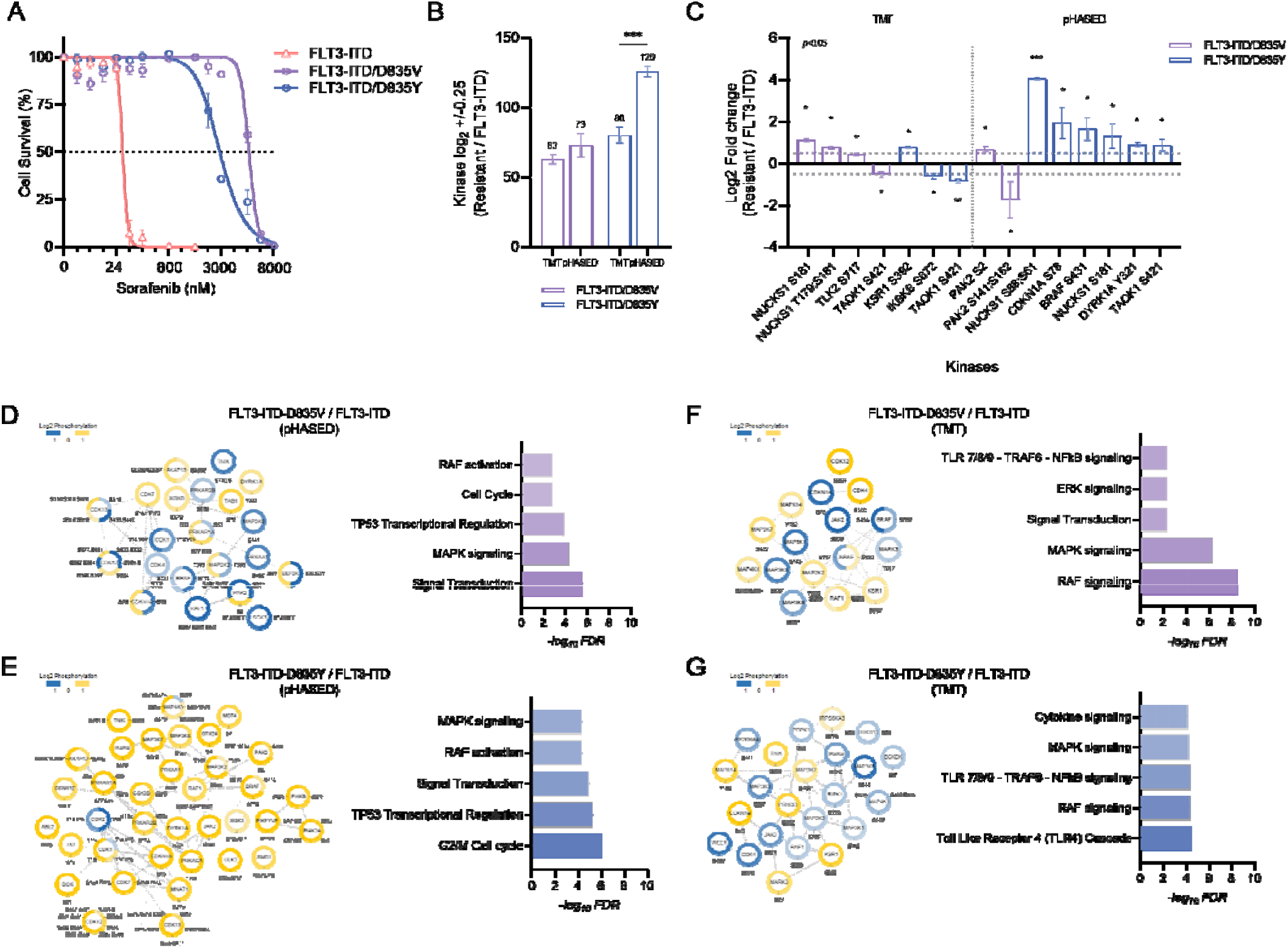
Comparison of phosphorylated kinases identified in resistant cell lines comparing TMT and pHASED experiments. *A*) Cell viability was assessed by resazurin assay at 48 h following treatment with sorafenib in FLT3-ITD, FLT3-ITD/D835V, and FLT3-ITD/D835Y isogenic cell lines (n=3 independent replicates). *B*) Number of kinases identified as differentially phosphorylated (log_2_ ± 0.25) in double mutants in comparison to FLT3-ITD cell using TMT and pHASED datasets. *C*) Kinases with significantly increased or decreased phosphorylation (log_2_ ± 0.25; p<0.05) in resistant cell lines comparison to FLT3-ITD identified by TMT and pHASED. Functional protein-protein interaction network and enrichment profile of kinases differentially phosphorylated (log_2_ ± 0.25) in resistant cell lines compared to FLT3-ITD cells. Protein interaction network shown corresponding to the major cluster of kinases identified in *D*) FLT3-ITD/D835V via pHASED; *E*) FLT3-ITD/D835V via TMT; and *F*) FLT3-ITD/D835Y via pHASED, and *G*) FLT3-ITD/D835Y via TMT. Yellow indicates increased phosphorylation, whereas Blue represents decreased phosphorylation. Canonical pathways of enriched kinases (Purple = FLT3-ITD/D835V, Blue = FLT3-ITD/D835Y) with FDR <1% in each comparison are also shown.

Protein-protein interaction network analysis of kinases identified in TMT and pHASED (log2 +/-0.25) revealed the enrichment of clustered kinases associated with signaling pathways that are known to be commonly deregulated in cancer (Fig. 5*D-G*). Both datasets identified kinases associated with, Signal Transduction, RAF (40) and ERK/MAPK (41) signaling (Fig. 5*D-G*), in line with previous studies that show potent activation of this oncogenic signaling pathway drives resistance to sorafenib (42). In addition, across resistance models, pHASED identified kinases responsible for controlling cell cycle and p53 signaling (43) (Fig. 5*D*, 5*E*), whereas TMT identified kinases associated with NFkB signaling (44) (Fig 5*F*, 5*G*).

### pHASED identified divergent DNA damage and repair pathways associated with sorafenib resistance in FLT3-mutant AML

Kinase-substrate enrichment analysis (KSEA) of TMT and pHASED datasets comparing resistant to diagnosis cells, predicted activation of oncogene RAC-alpha serine/threonine-protein kinase (AKT1), cell cycle regulators Cyclin dependent kinase 16 (CDK16), Polo-like kinase 2/3 (PLK2, PLK3), Serine/threonine-protein kinase VRK1/2 (VRK1/2), and Serine/threonine-protein kinase Kist (UHMK1); and DNA damage sensor and repair associated DNA-dependent protein kinase (PRKDC, or DNA-PK) (Fig. 6*A,* Supplemental Tables S*10*, S*11*). Furthermore, IPA analysis of canonical pathways associated with resistance (Fig. 6*B,* Supplemental Tables S*12*, S*13*) uncovered cell cycle regulation (Cell cycle control of Chromosomal Replication, G1/S and G2/M checkpoints, and Cyclins and Cell Cycle Regulation), and DNA damage and repair (ATM, NER, p53 and BRCA1, *p*-value <0.001) signaling as among the most significant enriched canonical pathways in resistant cell lines. Both mutants were also enriched for FLT3, and AML associated signaling pathways such as ERK/MAPK, JAK/STAT, mTOR, and PI3K/AKT, although with less statistical power (increased *p*-value ≤0.05).

**Figure 6.**
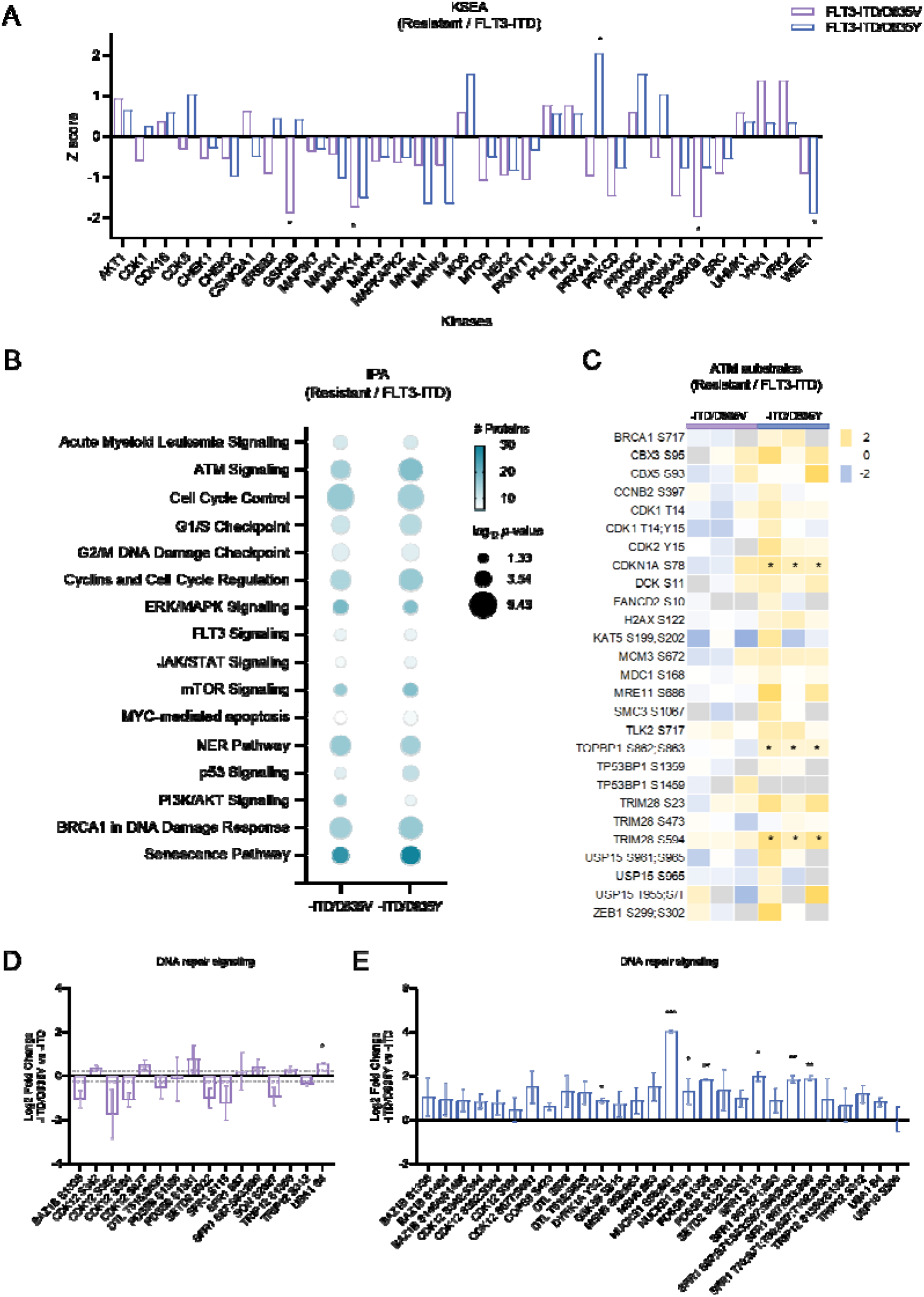
Bioinformatic analysis of FLT3 resistant phenotype via pHASED. *A*) Kinase substrate enrichment analysis (KSEA) profile of resistant cell lines compared to FLT3-ITD. Z score indicates predicted kinase activity, with a positive value predictive of kinase activation and a negative value predictive of kinase inhibition. *B)* Ingenuity Pathway analysis (IPA) of phosphorylated changes in resistant cell lines compared with FLT3-ITD. *C*) Phosphorylation profile of ATM substrates in resistant cell lines compared with FLT3-ITD. Yellow indicates increased phosphorylation, whereas Blue represents decreased phosphorylation. Missing values are colored Grey. Phosphorylation profile of substrates downstream DSB repair pathway for *D*) FLT3-ITD/D835V and *E*) FLT3-ITD/D835Y compared with FLT3-ITD. Statistical significance calculated via one-way ANOVA with significant threshold of \**p*<0.05, \*\**p*<0.01 and \*\*\**p*<0.001.

The Serine/threonine protein kinase (ATM) regulates response to DNA damage caused by double-strand breaks (DSBs) (13). ATM is member of the phosphoinositide 3-kinase (PI3K)-related protein kinase (PIKK) family, and signals through DNA damage response kinases ATR, DNA-PKcs and Nonsense Mediated MRNA Decay Associated PI3K Related Kinase (SMG1) (45, 46). One potent mechanism of increased DSBs is via the excess production of reactive oxygen species (ROS). Increased ROS production by the NADPH oxidase (NOX) family of enzymes in acute leukemias, particularly FLT3-ITD AML, has been increasingly studied over the last few years, and highlights that elevated ROS is a mechanism conferring survival advantages in FLT3-mutant AML (47–50). Given ATM signaling was predicted to be one of the top ranked canonical pathways driving the DNA damage repair and response pathways in resistant cells (Fig. 6*B*), we chose to analyze this pathway to test the biological utility of the phosphoproteomic analysis generated via pHASED in FLT3-ITD/D835V and FLT3-ITD/D835Y mutant cells.

Analysis of ATM (Fig. 6*C*) and DSB repair phosphoproteins (Fig 6*D*, 6*E*) in resistant models using pHASED, revealed divergent phosphorylation profiles. In FLT3-ITD/D835V mutant cells, pHASED only identified significantly increased phosphorylation of DSB repair pathway phosphoprotein UBA1 (S4) (log2 0.59; *p*=0.037) (Fig. 6*D*). Whereas, in FLT3-ITD/D835Y mutant cells, pHASED identified significantly increased phosphorylation of phosphoproteins downstream of ATM kinase signaling including CDKN1A (S78) (log2 1.62; *p*=0.034), TOPBP1 (S862, S863) (log2 1.00; *p*=0.048) and TRIM28 (S594) (log2 2.06; *p*=0.04) (Fig. 6*C*). Additionally, increased phosphorylation of three phosphopeptides for SFR1 (S67, S83, S99; S67, S71, S83, S87, S99, S103; and S115) (log2 1.93, *p*=0.003; log2 1.88 *p*=0.010; and log2 2.02 *p*=0.028, respectively), DYRK1A (Y321) (log2 0.88; *p*=0.03), two phosphopeptides for NUCKS1 (S58, S61; and S181) (log2 4.07, *p*=0.0009; and log2 1.28, *p*=0.04, respectively), and PDS5B (S1356) (log2 1.84; *p*=0.022) were identified in FLT3-ITD/D835Y mutant cells, highlighting the unique mechanisms of DNA repair regulation in this resistance model (Fig. 6*E*). These data add further evidence to pathway analysis divergencies shown using KSEA (Fig. 6*A*) and IPA canonical pathways association analysis comparing the double mutants (Fig. 6*A*).

### Combined inhibition of ATM and FLT3 showed synergistic effect in sorafenib-resistant cell lines

Combination cytotoxicity analysis using the ATM inhibitor KU-60019 (29) in combination with the FLT3 inhibitor sorafenib (28) was highly synergistic, particularly in cells harboring the sorafenib resistance mutation FLT3-ITD/D835Y (Fig. 7). In accordance with pathway prediction bioinformatic analyses (Fig. 6), FLT3-ITD/D835Y mutant cells showed increased sensitivity to the combination, resensitizing cells to sorafenib (Fig. 7*A*, Bliss synergy analysis score 12.42; 0.062 µM sorafenib, 1.25 µM KU-60019); whereas FLT3-ITD/D835V mutant cells showed maximal synergy at higher doses (Fig. 7*B*, Bliss score 10.71; 500 nM sorafenib, 2.5 µM KU-60019). The combined inhibition of ATM and FLT3 signaling was only additive in FLT3-ITD mutant cells (Bliss score 5.20) with these cells highly sensitive to sorafenib alone (Fig. 5*A*, 7*C*). To test the *in vitro* preclinical benefits using physiological concentrations of sorafenib, cell survival comparisons were performed at 0.062 µM sorafenib. Again, these data confirmed the increased synergistic effects of combined ATM and FLT3 inhibition in FLT3-ITD/D835Y mutant cells (Fig. 7*D*) compared with FLT3-ITD/D835V mutant cells (Fig. 7*E*). Together, these results confirm that ATM inhibition plays a role in the resensitization of FLT3-ITD/D835 mutant resistant cells to sorafenib, validating the pHASED phosphoproteomic prediction of the important role ATM signaling plays in signaling downstream of the FLT3-ITD/D835Y mutation.

**Figure 7.**
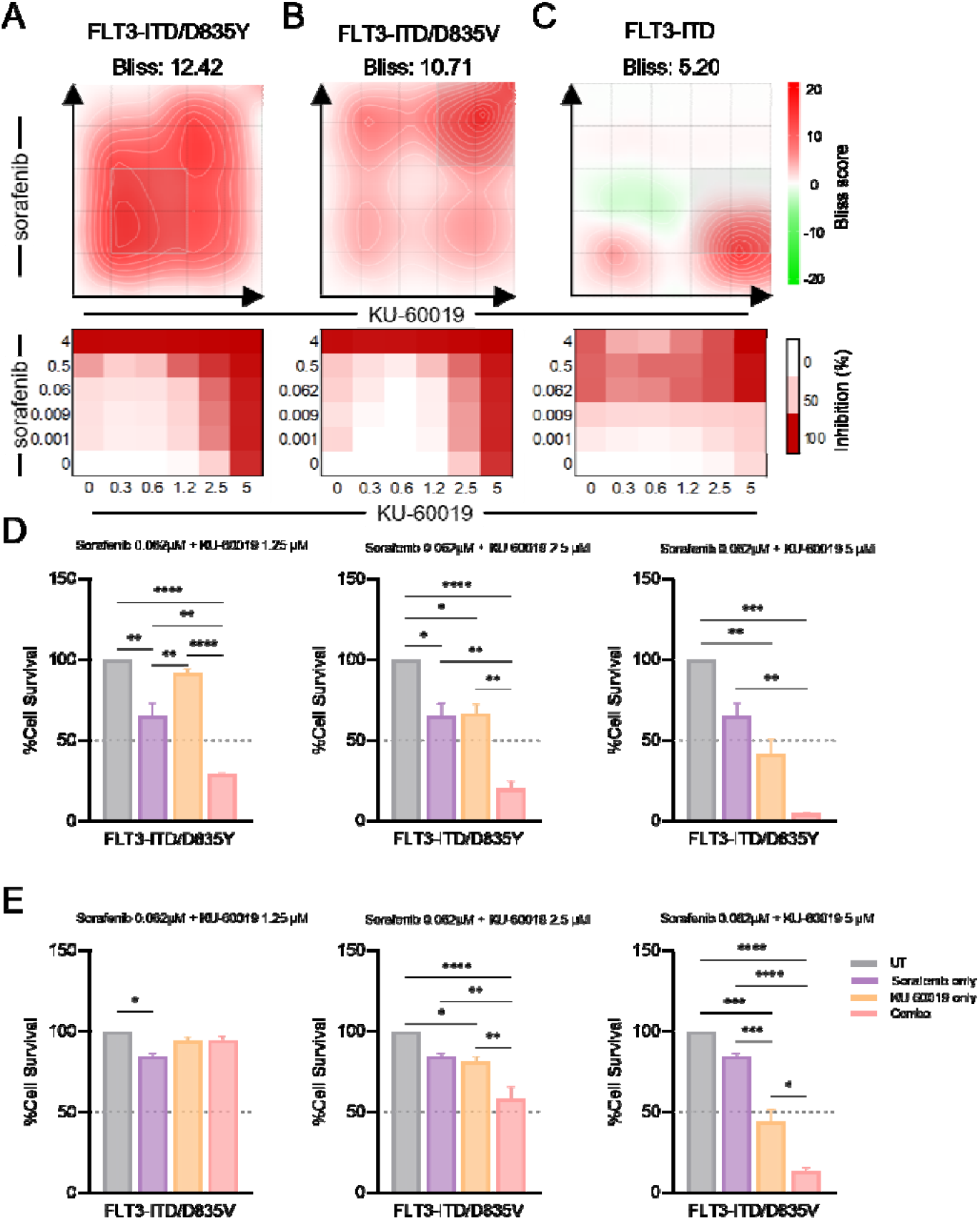
Sensitivity to ATM inhibition (KU-60019) in combination with FLT3 inhibitor sorafenib. Bliss synergy analysis of combined effect of sorafenib and KU-60019 in *A)* FLT3-ITD/D835Y, *B)* FLT3-ITD/D835V, and *C)* FLT3-ITD (<0= antagonistic, >0<10=additive, >10=synergistic). Cell survival comparison of *D*) FLT3-ITD/D835Y, and *E*) FLT3-ITD/D835V cell lines at 62 nM sorafenib in combination with 1.25 µM, 2.5 µM and 5 µM KU-60019 (n=3 independent replicates). Statistical significance calculated via one-way ANOVA with significant threshold of \**p*<0.05, \*\**p*<0.01 and \*\*\**p*<0.001.

## DISCUSSION

Proteomics and phosphoproteomics have been acknowledged as being among the most effective strategies to predict drug sensitivities (1, 51). However, we are yet to establish phosphoproteomic profiling in the clinical setting, or even to provide such as an additive resource to genomically predicted therapeutic strategies; the establishment of which would represent a pivotal advance in precision-medicine treatment regimens. Clinical phosphoproteomic profiling also has enormous potential to identify treatment targets that are invisible to genomics approaches, or to be used as an indicator of prognosis in the *de novo* and refractory settings, in real-time. Indeed, the optimization of pHASED reported herein, goes some way to moving phosphoproteomics from the discovery laboratory to that of the well-equipped pathologist. Importantly, the reduced complexity and sample preparation time of pHASED provides users with the capacity to prepare and sequence the phosphoproteomics of any biological system in less than a week. Furthermore, pHASED provided accurate LFQ and online deconvolution using FAIMS, whilst maintaining deep phosphoproteomic coverage without the need for offline 2D-LC techniques.

FAIMS was initially and elegantly optimized to provide single-shot LC-MS/MS results that compared favorably with 2D-LC fractionation experiments (22). Specifically, FAIMS was first reported in the context of analyzing the non-modified proteome of a cell line established from a chronic myelogenous leukemia patient (K562). In this study, the use of six CVs during a six-hour single-shot FAIMS experiment identified 8,007 non-modified proteins; comparable to the 7,776 non-modified proteins identified by the use of four 2D-LC fractionated samples, each analyzed for 1.5 h. Here, we have optimized 5 h of FAIMS using four CVs identifying 1,587 quantified phosphoproteins using pHASED, compared to nine 2D-LC fractions sequenced over ∼19 h, which identified and quantified 1,958 phosphoproteins using a TMT approach.

Although TMT identified more unique phosphoproteins than pHASED, it is well established that the use of isobaric tags can compromise identification efficiency due to peptide ratio compression artifacts caused by coeluting ions alongside that of the precursor ion of the peptide of interest; a phenomenon that interferes with MS2 based reporter ion quantitation (19, 52). Hence, this can limit the dynamic range of quantitation, and can often underrepresent the biological variability that exists between samples, especially for low abundant proteins (11, 53). Furthermore, the use of isobaric tags alters charge states during electrospray ionization (54), with the cleavage and loss of isobaric tags during MS2 generating fragment ions that complicate spectral interpretation by database searching algorithms (55), contributing to reduced identification efficiency. This was again evidenced by comparing phosphopeptide changes in FLT3-mutant AML models (n=5) with that of isogenic cell lines transduced to express the wt-FLT3 receptor (n=1), where the log2 fold-change of the TMT experiment was 0, compared with pHASED which showed an average log2 fold-change of 1.29 (*p*=0.0002). The biological differences revealed by pHASED further highlight the observation that knock in of each of the FLT3-mutations induced autonomous growth of isogenic cell lines, whereas cell lines transduced to express the wt-FLT3 receptor, required supplementation of growth factors to maintain growth and survival (5, 6, 56).

It was also of interest to note that pHASED identified more unique phosphosites per phosphoprotein compared to TMT. The biological context of this result was investigated by analyzing signaling pathways identified by both MS approaches to determine whether the increased number of identified phosphosites provided molecular insights relevant to the dissection of therapeutic vulnerabilities. Indeed, using both TMT and pHASED, IPA predicted ATM signaling to show increased activity in both FLT3-ITD/D835V and FLT3-ITD/D835Y double mutant cell line models compared to cells harboring FLT3-ITD mutations alone. ATM plays a functional role in the cellular response to DNA DSBs. Here it protects the cell against genotoxic stress, but, in cancer cells, helps to drive resistance to anticancer therapies thus favoring leukemic growth and survival (57). Therefore, it is unsurprising that ATM signaling may play a role in resistance to sorafenib in cells harboring double mutant FLT3-ITD/D835. However, sorafenib is not only a potent inhibitor of wt-FLT3 and FLT3-ITD, but also inhibits other receptor tyrosine kinases including VEGFR, PDGFR, KIT and RET, as well as downstream serine/threonine kinases including RAF/MEK/ERK (28). Indeed, in sorafenib-resistant double mutant FLT3-ITD/D835 cells, significantly increased MEK/ERK signaling was predicted when compared with the phosphoproteomes of FLT3-ITD mutant cells, thereby helping to explain the 47-fold increase in IC_50_ seen between the cell types. In glioma cells, ATM inhibitors increased radiotherapy sensitivity (29, 58, 59), with ATM signaling through the RAF/MEK/ERK pathway critical for radiation-induced ATM activation, suggestive of a regulatory feedback loop between ERK and ATM (60). Sorafenib dose-dependently induced the generation of ROS in tumor cells *in vitro* and *in vivo* (61), and hence it is highly possible that in FLT3-ITD/D835 double mutants, RAF/MEK/ERK signaling through ATM helps to maintain proliferation and promote DNA repair, even under situations of genotoxic stress induced by high dose sorafenib.

pHASED identified more significant phosphorylation changes in ATM substrates in FLT3-ITD/D835Y cells compared to FLT3-ITD/D835V cells (*p*≤0.05). Combination cytotoxicity assays revealed significantly increased synergy between sorafenib and the ATM inhibitor KU-60019 at physiologically relevant doses (most strikingly in FLT3-ITD/D835Y cells) providing a treatment paradigm for patients harboring sorafenib resistance. The increased phosphosite coverage arising from pHASED analyses potentially provides a more accurate indication of the regulation of the ATM signaling pathway, and hence highlights mechanisms promoting resistance to sorafenib (12); information that can be exploited to tailor effective preclinical treatment strategies.

Although pHASED may afford the opportunity to perform an unrestricted number of analyses (of benefit in the clinical setting where cancer diagnosis may not follow a predictable schedule), there remains important questions on how phosphoproteomics would be practically implemented as a clinical decision-making tool. For example, consideration needs to be given to sample processing time and the methods of patient sample collection; the phosphoproteome of leukemic blasts isolated from the bone marrow will differ from that sequenced from leukemic blasts isolated from peripheral blood. Additionally, the steps taken to enrich leukemic blasts following bone marrow trephine biopsy or phlebotomy are to be considered as alterations in signaling pathway activity can be influenced simply by the culture media used, or even the type of blood tube used at the time of sample collection (25), necessitating optimization and standardization of workflows. Importantly, for phosphoproteomics to aid in the treatment of cancer, the assessment of which pathways should be targeted and by which drugs needs to be evaluated under clinical trial conditions, like those testing whole genome sequencing (WGS) and RNA sequencing (RNAseq) strategies, in order to ensure robust recommendations can be made given phospho/proteomic data generated via pHASED (62).

In summary, the data obtained in the present study provides a novel method for LFQ of high-throughput phosphoproteomic data that maintains deep phosphoproteomic coverage without the need for complex 2D-LC strategies. pHASED provides the flexibility to analyze samples as they present and is not limited by the number of analyses that can be performed. Reduced time and complexity in sample preparation, and the optimization of online phosphoproteome deconvolution using a stepped CV FAIMS interface, provided accurate and reproducible phosphoproteomes of complex cancer cells in less than a week. Moreover, pHASED successfully identified novel drug targets and potential therapeutic strategies to treat AML models resistant to therapies used in the clinic; optimized technologies that we hope will help in the rapid characterization of highly aggressive forms of cancer, as an important step towards improving treatment outcomes for cancer sufferers.

## ACKNOWLEDGMENTS

This study was supported by Cancer Institute NSW Fellowships (M.D.D., N.MV). M.D.D. is supported by an NHMRC Investigator Grant – GNT1173892. This project is supported by an NHMRC Ideas Grant APP1188400 and NHMRC Targeted Research Grant GA65801. The contents of the published material are solely the responsibility of the research institutions involved or individual authors and do not reflect the views of NHMRC. D.S. and T.M. are supported by Zebra Equities Ph.D. Scholarships. Grants from the Hunter Medical Research Institute, Hunter Children’s Research Foundation, Jurox Animal Health, Zebra Equities, Hunter District Hunting Club and Ski for Kids, and The Estate of James Scott Lawrie supported this work. The Cancer Institute NSW in partnership with the Faculty of Health and Medicine from the University of Newcastle funded the MS platform. Figure 1 was created with BioRender.com

## AUTHOR CONTRIBUTIONS

Contribution: D.E.S., and M.D.D., conceived and designed the study and interpreted the results. D.E.S., H.C.M, D.A.S.B., N.D.S., M.F.J., R.S.K., R.J.D., Z.P.G., T.M., E.R.J., I.J.F., P.S.K., A.M., H.P.M., and M.D.D., conducted the experiments and performed data analysis. A.M.D., B.N., and N.M.V., provided discipline specific expertise; D.E.S., and M.D.D., wrote and edited the manuscript. All authors discussed the results and commented on the manuscript.

## DATA AVAILABILITY

The mass spectrometry proteomics data have been deposited to the ProteomeXchange Consortium (http://proteomecentral.proteomexchange.org) via the PRIDE partner repository (63) with the dataset identifier PXD032204 (TMT); and PXD032296 (pHASED).

PRIDE Reviewer account details:

TMT:

Username

Password: tGkAKsrp

pHASED:

Username

Password: 9yGk4Lrt

## ABBREVIATIONS

ACN: acetonitrile
AKT1: RAC-alpha serine/threonine-protein kinase
AML: acute myeloid leukemia
ATM: Serine/threonine protein kinase ataxia-telangiectasia mutated
ATR: Serine/threonine-protein kinase ataxia- and Rad3-related
AURKB: Aurora kinase B
BCA: bicinchoninic acid
CSNK1D: Casein kinase I isoform delta
CSNK2A2: Casein kinase II subunit alpha
CV: compensation voltage
DDA: data dependent acquisition
DSB: double-strand breaks
DTT: dithiothreitol
FAIMS: high-field asymmetric waveform ion mobility spectrometry
FDR: false discovery rate
HCD: high-energy collision dissociation
HCl: hydrochloric acid
HILIC: hydrophilic interaction liquid chromatography
IPA: ingenuity pathway analysis
ITD: internal tandem duplications
KSEA: kinase-substrate enrichment analysis
LC: liquid chromatography
LFQ: label-free quantitation
m/z: mass-to-charge ratios
MS: mass spectrometry
nLC-MS/MS: nanoliquid chromatography–tandem mass spectrometry
NM: non-modified
NOX: NADPH oxidase
pHASED: Phospho Heavy-labeled-spiketide FAIMS Stepped-CV DDA
PI3K: phosphoinositide 3-kinase
PIKK: Phosphatidylinositol 3-kinase-related kinases (PIKKs)
PRKDC; DNA-PK: DNA-dependent protein kinase
PSM: phosphopeptide-spectrum matches
PTMs: posttranslational modifications
ROS: reactive oxygen species
RT: room temperature
SMG1: Nonsense mediated mRNA decay associated PI3K related kinase
SPE: solid phase extraction
TCA: trichloroacetic acid
TEAB: triethylammonium bicarbonate
TFA: trifluoroacetic acid
TiO2: titanium dioxide
TKI: tyrosine kinase inhibitors
TMT: tandem mass tag
UHMK1: Serine/threonine-protein kinase Kist

