## Supporting Information for "High-Throughput Global Phosphoproteomic Profiling Using Phospho Heavy-Labeled-Spiketide FAIMS Stepped-CV DDA (pHASED)"

**SUPPLEMENTARY MATERIALS AND METHODS**

*Plasmid and Vector Construction*

FLT3 wild-type (FLT3-WT) and FLT3 mutant constructs were cloned into murine stem cell virus-green fluorescent protein (pMSCV-GFP) vectors, of which templates were a kind gift from Leonie K. Ashman (University of Newcastle, Australia). Internal tandem duplication mutation was generated through the insertion of a 69bp fragment in FLT3 (FLT3-ITD). Mutagenesis was performed using QuikChange II Site-Directed Mutagenesis Kit (Agilent) as per manufacturer’s instructions, to generate FLT3-D835V and FLT3-D835Y mutants with a substitution of 3bp in codon D835 (GAT). FLT3-D835V mutant constructs were generated using “GGATTGGCTCGAGTTATCATGAGTGATTCC” and “GGAATCACTCATGATAACTCGAGCCAATCC” primers, whereas “GGATTGGCTCGATATATCATGAGTGATTCC” and “GGAATCACTCATGATATATCGAGCCAATCC” primers were used to generate FLT3-D835Y mutants (Sigma). FLT3-ITD/D835V and FLT3-ITD/D835Y double mutant receptors were generated with both -ITD and -D835 mutations. Vector mutations are listed in Table S14.

*Retroviral transduction*

Viral vectors were transformed using XL-Blue Supercompetent cells (Agilent) and extracted DNA (8µg) transfected into Phoenix Eco cells (Leonie K. Ashman,University of Newcastle). FLT3 mutant myeloblasts were generated through retroviral transduction on FDC-P1 cells using pMSCV-GFP vectors containing different FLT3 wild-type (FLT3-WT) and mutant cDNAs or empty vector (EV). Cells were selected using puromycin and transformation efficiency analyzed by Sanger DNA sequencing. Successfully transduced FDC-P1 cells were sorted to generate homogeneous populations of each cell type using GFP protein expression (>80%).

*Sanger Sequencing*

DNA was extracted from FDC-P1 transduced cell lines using the Wizard SV Genomic DNA Purification System (Promega), as per manufacturer’s instructions and quantified using NanoDrop 2000 (Thermo Scientific). 50ng of DNA were separated for Polymerase chain reaction (PCR) amplification and ran in 1% agarose gel at 100V. DNA was extracted from gel using the Wizard SV Gel and PCR Clean-up System (Promega), as per manufacturer’s instructions, and sent for Sanger sequencing by the Australian Genome Research Facility (AGRF, Australia).

**SUPPLEMENTARY FIGURE**

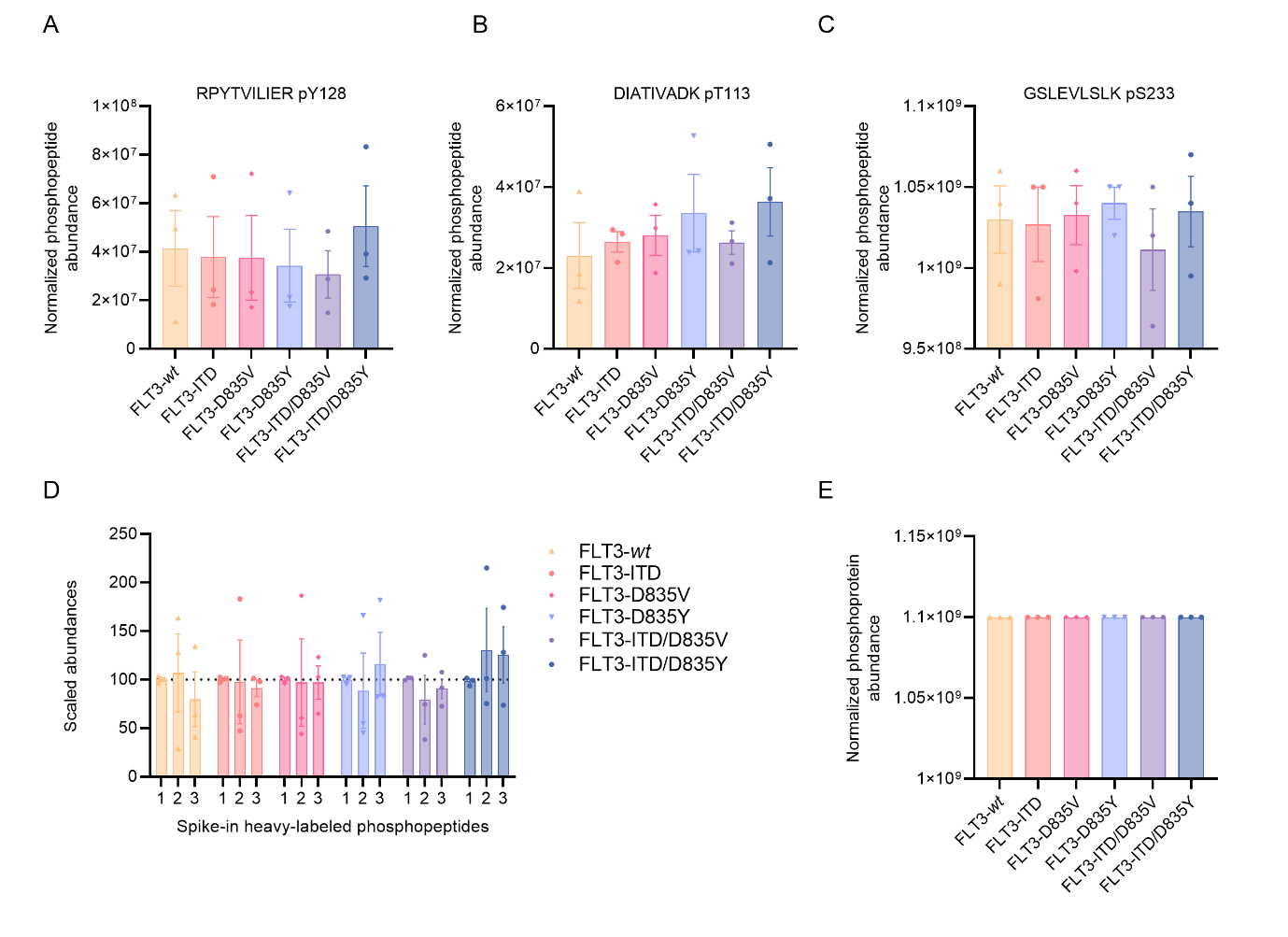

**Supplementary Figure S1**. **Standard spike-in control abundances (n=3 biological replicates).** Normalized abundances for heavy-labeled spike-in peptides phosphorylated at *A*) tyrosine, *B*) threonine, and *C*) serine residues for FLT3-wt and FLT3-mutant cell lines. *D*) Scaled abundances for the three spike-in heavy-labeled phosphopeptides used as internal controls. *E*) Normalized phosphoprotein abundance identified based on the three heavy-labeled phosphopeptide controls.
